# Tractography in the presence of white matter lesions in multiple sclerosis

**DOI:** 10.1101/559708

**Authors:** Ilona Lipp, Greg D Parker, Emma Tallantyre, Alex Goodall, Steluta Grama, Eleonora Patitucci, Phoebe Heveron, Valentina Tomassini, Derek K Jones

**Author notes:** shared first authorship (Ilona Lipp, Greg D Parker).

## Abstract

Accurate anatomical localisation of specific white matter tracts and the quantification of their tract-specific microstructural damage in multiple sclerosis (MS) can contribute to a better understanding of symptomatology, disease progression and intervention effects. Diffusion MRI-based tractography is being used increasingly to segment white matter tracts as regions-of-interest for subsequent quantitative analysis. Since MS lesions can interrupt the tractography algorithms tract reconstruction, clinical studies frequently resort to atlas-based approaches, which are convenient but ignorant to individual variability in tract size and shape. Here, we revisit the problem of individual tractography in MS, comparing tractography algorithms using: (i) The diffusion tensor framework; (ii) constrained spherical deconvoution (CSD); and (iii) damped Richardson-Lucy (dRL) deconvolution. Firstly, using simulated and in vivo data from 29 MS patients and 19 healthy controls, we show that the three tracking algorithms respond differentially to MS pathology. While the tensor-based approach is unable to deal with crossing fibres, CSD produces spurious streamlines, in particular in tissue with high fibre loss and low diffusion anisotropy. With dRL, streamlines are increasingly interrupted in pathological tissue. Secondly, we demonstrate that despite the effects of lesion on the fibre orientation reconstruction algorithms, fibre tracking algorithms are still able to segment tracts that pass areas with high prevalence of lesions. Combining dRL-based tractography with an automated tract segmentation tool on data from 131 MS patients, the corticospinal tracts and arcuate fasciculi were successfully reconstructed in more than 90% of individuals. Comparing tract-specific microstructural parameters (fractional anisotropy, radial diffusivity and magnetisation transfer ratio) in individually segmented tracts to those from a tract probability map, we showed that there is no systematic disease-related bias in the individually reconstructed tracts, suggesting that lesions and otherwise damaged parts are not systematically omitted during tractography. Thirdly, we demonstrate modest anatomical correspondence between the individual and tract probability-based approach, with a spatial overlap between 35 and 55%. Correlations between tract-averaged microstructural parameters in individually segmented tracts and the probability-map approach ranged between *r* = .52 (*p* < .001) for radial diffusivity in the right cortico-spinal tract and *r* = .97 (*p* < .001) for magnetization transfer ratio in the arcuate fasciculi. Our results show that MS white matter lesions impact fibre orientation reconstructions but this does not appear to hinder the ability to anatomically localise white matter tracts in MS. Individual tract segmentation in MS is feasible on a large scale and could prove a powerful tool for investigating diagnostic and prognostic markers.

## 1. Introduction

Accurate quantification of white matter damage in multiple sclerosis (MS) is important for characterising disease pathophysiology (Barkhof et al., 2009). While the hallmark of pathology in MS is focal demyelinating lesions, correlations between the total volume of lesional tissue and disability are low, presenting the so-called clinical-radiological paradox (Barkhof, 1999, 2002). Disability and prognosis may be better explained by the anatomical location of lesions and of diffuse microstructural damage outside of lesions (Charil et al., 2003; Kolind et al., 2012). Accurate assignment of damage to specific anatomical white matter tracts can provide a more accurate picture of the disease (Lin et al., 2005), and is important for longitudinal studies that explore the effect of rehabilitation and experimental interventions on white matter in relevant tracts (Bonzano et al., 2014).

Anatomical localisation of white matter tracts in vivo currently relies on diffusion-weighted MRI and fibre tracking (Catani et al., 2002; Jeurissen and Leemans, 2017). The reconstruction of individual tracts is based on combining prior anatomical knowledge of the tract location with diffusion MRI-based evidence about fibre orientations in the imaging voxels. Early tractography work was performed by producing streamlines that follow the principal eigenvector of the diffusion tensor (e.g. Basser et al. (2000); Mori et al. (1999)), while more recent work relies on the estimation of the fibre orientation distribution function (e.g. Tournier et al. (2004)). Tractography can be combined with an extraction of tract-specific microstructural metrics (Jones et al., 2005, 2006) to be used for investigating individual differences or longitudinal changes in tract-specific microstructure (e.g. Lin et al. (2005); Metzler-Baddeley et al. (2011)).

Focal brain pathology, such as is present in MS, can affect tractography. MS lesions are characterised by fibre loss and consequent increase in extracellular water associated with tissue destruction, which is reflected in the diffusion profile (Filippi et al., 2001). During tractography, at each step of streamline reconstruction, angle and amplitude criteria are in place to avoid spurious tracking. Early tractography studies used fractional anisotropy (FA) as a criterion for streamline termination. As FA is significantly decreased in white matter lesions in MS (Filippi et al., 2001), applying tractography to data from MS patient has been recognized as being problematic (Ciccarelli et al., 2008; Inglese and Bester, 2010). In the absence of a strategy to the problem of streamline termination (such as employed by Lagana et al. (2011), Tench et al. (2002) and Wang et al. (2018)), the reconstructed tracts may lack anatomical accuracy, e.g. by premature termination of tracking (Ozturk et al., 2010). For this reason few studies apply tractography in patients (other examples are Lin et al. (2005); Reich et al. (2007, 2010)).

More recent advances in tractography algorithms do not rely on the estimated fibre orientation from the diffusion tensor (which yields one fibre orientation estimate per voxel), but employ deconvolution approaches that yield multiple fibre orientations per voxel (e.g. Tournier et al. (2004)). This advanced approach could permit the reconstruction of streamlines through MS lesions, given that enough fibres are present to generate a peak in the estimated fibre orientation distribution (FOD). However, it is possible that due to the fibre loss in lesions, the peak amplitudes may fall below the threshold normally used for termination of tracking. On the other hand, lowering the FOD amplitude threshold could lead to the tract reconstruction following spurious peaks, such as those arising from noise, since lesions are characterised by an increased component of water with isotropic diffusion, which can compromise orientation estimates (Dell’Acqua et al., 2010). To our knowledge, tractography results obtained with spherical deconvolution approaches in MS patients have not been systematically assessed.

Due to the challenges related to tractography in pathological tissue, an alternative approach for obtaining tract-specific measures in MS was suggested (Pagani et al., 2005; Hua et al., 2008). First, tracts are reconstructed in the native space of brains of healthy volunteers and then normalised to a common reference space, where a tract-probability map is created. Then, the data from patients are aligned with the same common reference space. To get a tract-specific measure of damage, the probabilistic atlas is used to calculate a weighted average for the microstructural metric of interest.

While the probability-map based method has the advantage that individual tracts are only reconstructed in healthy brains, there are also considerable short-comings. The approach relies on inter-subject co-registration of the images by normalising them to a common reference space. This normalisation is generally performed based on structural images, e.g. T1-weighted images, on which white matter appears homogeneous (Hua et al., 2008; Pagani et al., 2005; Reich et al., 2010). The approach therefore implicitly assumes that white matter tract anatomy is consistent across individuals and in states of health and disease. However, this assumption is unlikely to hold. Tract location and shape vary even between healthy individuals (Wassermann et al., 2012), and in MS white matter atrophy is well described (Ge et al., 2001) and could affect some tracts more than others (Kezele et al., 2008). Applying a probability mask may yield measures of microstructural damage that are likely to include information from other tracts in the vicinity and may therefore not be anatomically precise. This imprecision could be the explanation for the low correlations between individual tractography and probability-based measures that have previously been reported for some tracts (Reich et al., 2010). Another explanation could be that the individual tractography omits damaged parts of the tract, leading to not-representative and biased estimates.

In this work, we reassess the feasibility of performing individual tractography in patients. Using simulations as well as in vivo data sets from patients, we compared the effect of MS pathology on the performance of three tractography algorithms: (i) DTI based tracking; (ii) Constrained spherical harmonic deconvolution (CSD)-based tracking and (iii) damped Richardson-Lucy-based tracking. Then, we reconstruct corticospinal tracts (CST) and arcuate fasciculi (Arc) in a large number of patients to evaluate the practical implications of tracking through white matter regions with a high prevalence of lesions.

## 2. Methods

### 2.1. Data acquisition

#### 2.1.1. Simulated data

We simulated diffusion data for a sequence comparable to our in vivo sequence (Section 2.1.3), with a conservative SNR estimate of 20:1 for non-diffusion weighted images (see Supplementary Section 1.1.2), using Camino’s *datasynth* (Hall and Alexander, 2009). We simulated data for a number of tissue substrates, all characterised by impermeable parallel cylinders with mean radius of 1 µm and standard deviation of 0.7 µm. To assess the effect of fibre loss and increased extracellular volume fraction, as seen in lesion pathology, we simulated substrates that differed in their intracellular volume fractions, by varying the number of cylinders placed in the substrate. We simulated substrates with a single fibre population and substrates with two fibre populations crossing at 90 degrees. Details on the implementation of the simulations are reported in Supplementary Section 1.1.1.

#### 2.1.2. Participants

In total, data from 135 right-handed MS patients, who took part in a large-scale imaging project (Lipp et al., 2017), contributed to this study. Diffusion data were available for 131 patients. A subset (29 patients) of the large cohort had been age-and gender- matched to 19 healthy controls, who underwent the same scanning protocol. For all analyses, for which pathological tissue was compared to healthy control tissue, only data from the two matched groups were considered. For analyses regarding in vivo tract segmentation, data from all 131 patients were considered. Data from the healthy controls were used for creating the tract probability maps as well as a shape model that was used for automatic tract segmentation for the larger cohort of patients, as described below.

The clinical and demographic characteristics of patients and controls are presented in Table 1.

**Table 1:**
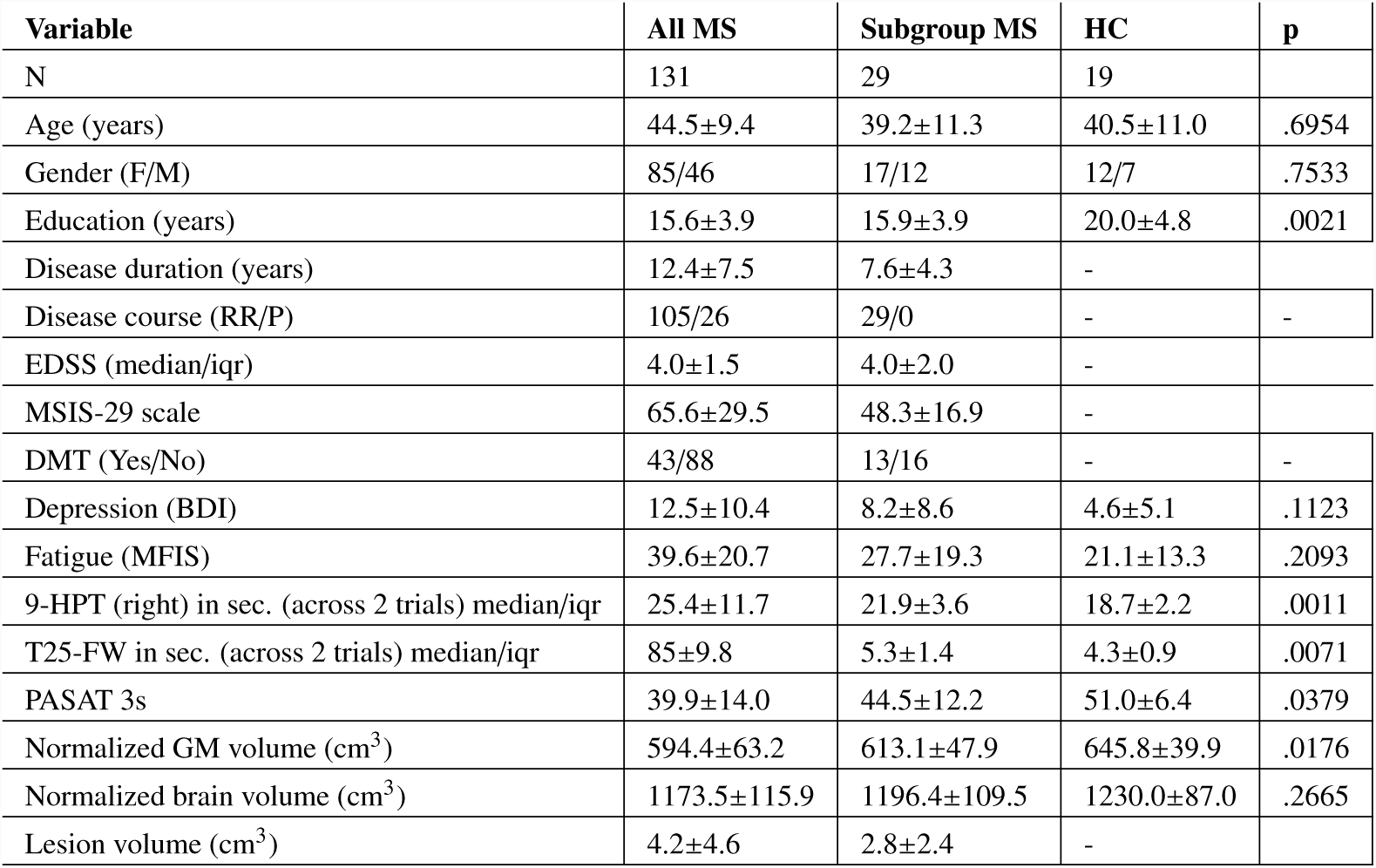
Demographic and clinical characteristics of the cohorts investigated. Characteristics are provided for all multiple sclerosis patients (MS), as well as the subgroup of patients matched to and healthy controls (HC). Unless otherwise indicated, descriptive statistics provided are means and standard deviations. For statistical comparison between the two matched groups (*p*-values are reported), Chi-square tests were computed for categorical variables, Kruskall-Wallis tests for skewed variables (9 hole peg test and timed 25 foot walk), and unpaired *t*-tests for the rest. P values for group differences are provided. **Acronyms:** RR = Relapsing-remitting, P = progressive MS (includes primary and secondary progressive patients), EDSS = Extended Disability Status Scale, MSIS-29 = Multiple Sclerosis Impact Scale 29 items, DMT = disease-modifying treatment, BDI = Beck Depression Inventory, MFI S= Modified Fatigue Impact Scale, 9-HPT: 9 hole peg test, T25-FW: timed 25 foot walk, PASAT = Paced Auditory Serial Addition Test (3 second version). Normalized brain and grey matter volume was calculated using SIENAX (Smith et al., 2002).

| Variable | All MS | Subgroup MS | HC | p |
| --- | --- | --- | --- | --- |
| N | 131 | 29 | 19 |  |
| Age (years) | 44.5±9.4 | 39.2±11.3 | 40.5±11.0 | .6954 |
| Gender (F/M) | 85/46 | 17/12 | 12/7 | .7533 |
| Education (years) | 15.6±3.9 | 15.9±3.9 | 20.0±4.8 | .0021 |
| Disease duration (years) | 12.4±7.5 | 7.6±4.3 | - |  |
| Disease course (RR/P) | 105/26 | 29/0 | - | - |
| EDSS (median/iqr) | 4.0±1.5 | 4.0±2.0 | - |  |
| MSIS-29 scale | 65.6±29.5 | 48.3±16.9 | - |  |
| DMT (Yes/No) | 43/88 | 13/16 | - | - |
| Depression (BDI) | 12.5±10.4 | 8.2±8.6 | 4.6±5.1 | .1123 |
| Fatigue (MFIS) | 39.6±20.7 | 27.7±19.3 | 21.1±13.3 | .2093 |
| 9-HPT (right) in sec. (across 2 trials) median/iqr | 25.4±11.7 | 21.9±3.6 | 18.7±2.2 | .0011 |
| T25-FW in sec. (across 2 trials) median/iqr | 85±9.8 | 5.3±1.4 | 4.3±0.9 | .0071 |
| PASAT 3s | 39.9±14.0 | 44.5±12.2 | 51.0±6.4 | .0379 |
| Normalized GM volume (cm<sup>3</sup>) | 594.4±63.2 | 613.1±47.9 | 645.8±39.9 | .0176 |
| Normalized brain volume (cm<sup>3</sup>) | 1173.5±115.9 | 1196.4±109.5 | 1230.0±87.0 | .2665 |
| Lesion volume (cm<sup>3</sup>) | 4.2±4.6 | 2.8±2.4 | - |  |

#### 2.1.3. MRI acquisition

In vivo MRI data were acquired on a 3T General Electric HDx MRI system (GE Medical Systems, Milwaukee, WI) using an eight channel receive-only head RF coil (GE Medical Devices). We acquired the following sequences: a T2/proton-density weighted and a fluid-attenuated inversion recovery (FLAIR) sequence for lesion identification and segmentation, a T1-weighted sequence for identification of T1-hypointense MS lesions and for registration, and a twice-refocussed diffusion-weighted sequence (40 uniformly distributed directions (Jones et al., 1999), b = 1200 s/mm^2^), and a 3D MT sequence. Latter was used to calculate an additional microstructural parameter independent of the diffusion-weighted images. The acquisition parameters of all scan sequences are reported in Supplementary Table 2.

**Table 2:** Comparison of the three fibre orientation reconstruction algorithms in vivo. For each algorithm (tensor, dRL, CSD) and each tissue type (*Ctrl, NAWM, T2L, T1l*), we calculated voxel-wise averages of the following parameters: the number of streamlines found in a voxel, the average FA / FOD amplitude across all streamlines found in a voxel, the number streamline terminations due to the amplitude threshold in a voxel, and the number streamline terminations due to the angle threshold in a voxel. The mean std of these measures across healthy controls (*Ctrl*; N = 19) and patients (*NAWM, T2L, T1L*; N = 29) are reported. The values across different tissues types were statistically compared (unpaired t-test between *Ctrl vs NAWM* tissue; paired t-tests for *T2L vs NAWM*, and *T1L vs T2L*) and *t* and *p* statistics are provided for each comparison. **Acronyms**: Ctrl: Control tissue, NAWM: normal appearing white matter, T2L: T2-weighted white matter hyperintense lesional tissue without T1-weighted hypointensity, T1L: T1-weighted white matter hypointense lesional tissue with corresponding T2-weighted hyperintensity.

|  | *Ctrl* | *NAWM* | *T2L* | *T1L* | *NAWM vs Ctrl* | *T2L vs NAWM* | *T1L vs T2L* |
| --- | --- | --- | --- | --- | --- | --- | --- |
| **Average number of streamlines per voxel** |  |  |  |  |  |  |  |
| Tensor | 32.31 ± 3.35 | 30.66 ± 4.23 | 33.48 ± 6.72 | 30.77 ± 9.52 | $t = -1.43, p = .16$ | $t = 2.64, p = .01$ | $t = -1.55, p = .13$ |
| dRL | 38.14 ± 5.91 | 39.40 ± 5.09 | 42.68 ± 8.61 | 40.41 ± 11.25 | $t = 0.79, p = .44$ | $t = 2.6, p = .02$ | $t = -1.3, p = .20$ |
| CSD | 63.37 ± 10.22 | 66.12 ± 8.77 | 70.70 ± 13.18 | 70.69 ± 18.86 | $t = 0.99, p = .33$ | $t = 2.4, p = .02$ | $t = -0.01, p = .996$ |
| **Average peak amplitude (FOD amplitude / FA) of all streamlines per voxel** |  |  |  |  |  |  |  |
| Tensor | 0.71 ± 0.03 | 0.68 ± 0.03 | 0.67 ± 0.04 | 0.64 ± 0.06 | $t = -3.1, p = .003$ | $t = -1.1, p = .29$ | $t = -2.4, p = .03$ |
| dRL | 0.22 ± 0.02 | 0.21 ± 0.02 | 0.21 ± 0.02 | 0.20 ± 0.03 | $t = -1.1, p = .29$ | $t = -0.2, p = .99$ | $t = -2.7, p = .01$ |
| CSD | 0.42 ± 0.04 | 0.43 ± 0.05 | 0.43 ± 0.06 | 0.42 ± 0.06 | $t = 0.78, p = .44$ | $t = 0.64, p = .53$ | $t = -2.2, p = .04$ |
| **Average number of streamline terminations due to amplitude threshold** |  |  |  |  |  |  |  |
| Tensor | 0.32 ± 0.01 | 0.24 ± 0.04 | 0.51 ± 0.26 | 0.97 ± 0.47 | $t = -8.7, p < .0001$ | $t = 5.8, p < .0001$ | $t = 8.5, p < .0001$ |
| dRL | 0.23 ± 0.01 | 0.25 ± 0.02 | 0.36 ± 0.11 | 0.70 ± 0.33 | $t = 5.9, p < .0001$ | $t = 5.2, p < .0001$ | $t = 6.9, p < .0001$ |
| CSD | 0.97 ± 0.03 | 0.87 ± 0.09 | 0.81 ± 0.19 | 0.80 ± 0.28 | $t = -4.7, p < .0001$ | $t = -1.7, p = .10$ | $t = -0.38, p = .71$ |
| **Average number of streamline terminations per voxel due to angle threshold** |  |  |  |  |  |  |  |
| Tensor | 0.10 ± 0.01 | 0.13 ± 0.02 | 0.16 ± 0.07 | 0.08 ± 0.07 | $t = 6.4, p < .0001$ | $t = 2.0, p = .05$ | $t = -6.4, p < .0001$ |
| dRL | 0.58 ± 0.02 | 0.71 ± 0.08 | 0.74 ± 0.20 | 0.53 ± 0.23 | $t = 7.2, p < .0001$ | $t = 0.86, p = .39$ | $t = -5.3, p < .0001$ |
| CSD | 1.37 ± 0.09 | 1.41 ± 0.12 | 1.76 ± 0.59 | 2.04 ± 0.76 | $t = 1.1, p = .26$ | $t = 3.2, p < .01$ | $t = 2.9, p = .0074$ |

### 2.2. Data analysis

#### 2.2.1. Algorithms for resolving fibre orientation

We compared three algorithms for recovering fibre orientational information. Firstly, as done in previous MS work (Hua et al., 2008; Reich et al., 2010), fibre orientation was estimated using the first eigenvector of the diffusion tensor. Additionally, we employed two fibre orientation distribution (FOD) deconvolution algorithms, which have been developed to overcome some of the limitations related to tensor-based tracking: a constrained spherical deconvolution (CSD, Tournier et al. (2007)), and a modified damped Richardson-Lucy algorithm (dRL, Dell’Acqua et al. (2010)). Deconvolution methods work by characterising a response function for a single fibre orientation. This response function is then deconvolved from the observed signal. The reason for considering both CSD and dRL is that previous work (Parker et al., 2013b) showed that while CSD performs better than dRL when resolving crossing fibres in voxels with low FA, it also more frequently produces spurious peaks and is more sensitive to mis-calibrations of the FOD. Details on the implementation of the two deconvolution algorithms are reported in Supplementary Section 1.1.3

#### 2.2.2. Analysis of simulated data

For each simulated voxel, we applied the three algorithms. From the resulting estimated fibre orientation profiles, we tested whether peaks could be correctly identified: a) along the true underlying direction(s) along which the cylinders had been placed; and b) along false direcions (the direction(s) orthogonal to the long axis of the simulated cylinders). For each substrate type and algorithm, the proportion of simulated voxels in which the reconstructed peak orientation closest to the simulated orientation subtended an angle of less than 45 degrees and reached the specified amplitude threshold (dRL: > 0.05, CSD: > 0.1, tensor: *FA* > .2) was determined. Additionally, the orientation dispersion of the detected peaks as a measure of algorithm precision was calculated using Basser et al. (2000)’s coherence measure to an average dyadic tensor which was calculated across all identified peaks (Jones, 2003). The dispersion measure can take values between 0 (all peaks point in exactly the same direction) and 1 (the detected peaks are uniformly distributed on the unit sphere).

#### 2.2.3. In vivo lesion mapping and segmentation

Damage was quantified in three tissue-states, which were expected to vary in their underlying microstructural damage: normal appearing white matter, T2-weighted white matter hyperintense lesional tissue without T1-weighted hypointensity, and T1-weighted white matter hypointense lesional tissue with corresponding T2-weighted hyperintensity, as reported elsewhere (Lipp et al., submitted). Briefly, normal appearing white matter was defined as FSL FAST (Zhang et al., 2001) segmented (80% thresholded) white matter at least 5 mm away from lesions. We classified lesional voxels as T1-weighted white matter hypointense if their signal intensity lay at least 1.5 interquartile ranges below the lower quartile of the distribution in normal appearing white matter. All other lesional voxels were classified as T2-weighted white matter hyperintense lesional tissue without T1-weighted hypointensity. Further, we restricted all three tissue classes to lesion-susceptible white matter (white matter with > 5% lesion probability, as defined by a lesion probability map derived from our data).

#### 2.2.4. MT processing

The MTR was calculated voxel by voxel with the equation MTR=[(S_0_-S_MT_)/S_0_]x100, whereby S_0_ represents the signal without the off-resonance pulse and S_MT_ represents the signal with the off-resonance pulse. The MTR images in native space were skull-stripped using FSL BET and non-linearly registered to the respective skull-stripped T1-weighted images using Elastix (Klein et al., 2010).

#### 2.2.5. Diffusion preprocessing

The DTI data were preprocessed in ExploreDTI (v 4.8.3; Leemans et al. (2009)). Data were corrected for head motion, distortions induced by eddy currents and EPI-induced geometrical distortions by registering each diffusion image to the corresponding T1-weighted anatomical image (Irfanoglu et al., 2012) using Elastix (Klein et al., 2010), with appropriate reorientation of the diffusion encoding vectors (Leemans and Jones, 2009). The anatomical image was first skull stripped and down-sampled to 1.5mm in order to reduce computation times during further processing of the diffusion data. RE-STORE (Chang et al., 2005) was used to account for outliers.

#### 2.2.6. Tractography algorithm

Whole brain tractography was performed in native space (for the subsample of 29 patients with all three algorithms, and for the entire sample of 131 patients using dRL), using an adaptation of CSD-based streamline tractography (Jeurissen et al., 2011; Tournier et al., 2004, 2007, 2008). Seed points were evenly distributed across vertices of a 2mm isotropic grid and propagated in 1mm steps with streamline length constraints of 20 −500mm. The diffusion tensor / fODF peaks were resolved at each new location (Jeurissen et al., 2011). In the case of CSD/dRL-based tracking, tracking was terminated if the fODF threshold fell below the defined threshold or the direction of streamline changed through an angle greater than 45°between successive steps. In the case of tensor-based tracking, instead of an FOD amplitude threshold, an FA threshold of .2 was used. The same procedure was then repeated by tracking in the opposite direction from the initial seed-points.

#### 2.2.7. Individual tract segmentation

For the purpose of this paper, the CST and arcuate fasciculi were segmented. The CST originates from motor and premotor cortices and runs to midbrain and medulla, passing the corona radiata and internal capsule (Al Masri, 2011), whose periventricular spaces are common spots for lesions (Kincses, 2010). The arcuate fasciculi connect the perisylvian cortex of the frontal, parietal, and temporal lobes (Catani and Thiebaut de Schotten, 2008).

Three-dimensional tractograms for specific white matter tracts were extracted from the whole-brain tractograms by applying multiple way-point of interest gates (Catani et al., 2002), drawn on color-coded fibre orientation maps (Pajevic and Pierpaoli, 1999). We applied tract reconstruction protocols to the data from the matched 29 patients and 19 healthy controls. All tract segmentations for a given tract were performed by the same operator (CST: EP, Arcuate fasciculus: SG). The protocol followed for segmentation is described in Supplementary Section 1.1.4. To assess inter-operator spatial agreement, another operator (SP) dissected all tracts in the data from the first five healthy controls, and spatial overlap Dice coefficient scores were calculated (see also Dice (1945); Zijdenbos et al. (1994); for details see Supplementary Section 1.1.5).

In the healthy control data, the approach of Parker et al. (2013a) was used to construct a shape model for each tract in each hemisphere from the manually segmented tracts. The resulting models were then used to automatically extract the tracts of interest across all datasets (patients and controls). All automatically segmented tracts were visually inspected and spurious streamlines were removed if necessary. To validate the automated protocol, for each of the 29 patients and 19 controls with both manually dissected and automatically dissected tracts, we calculated the spatial agreement between the two tract masks.

#### 2.2.8. Tract probability maps

For both patients and controls tract probability maps were computed, which indicate each voxel’s likelihood of being part of an individual’s tract. Each tract was first converted to a binary voxel-wise mask, indicating which voxels a tract intersected. To exclude voxels with only minimal streamlines, the 25% of voxels with lowest number of streamlines were ignored during this process. The exported tracts of each participant were registered to MNI space. This was done by first registering each participant’s structural high-resolution T1-weighted image to MNI space, using ANTs SyN (Avants et al., 2008) and then applying the warp to the tract NIfTI file (which had already been registered to the high resolution structural scan as part of the pre-processing pipeline). From the binary tract masks in MNI space, we computed probability maps for each tract and hemisphere. The probability maps for controls were then used to employ the probability-map based approach for obtaining tract-specific microstructural measures.

#### 2.2.9. Extraction of microstructural damage within tracts

For individual tracts, we computed the microstructural parameters (FA, RD, MTR) at each point along the individual streamlines (Jones et al., 2005) by trilinear interpolation of the surrounding voxels. We then computed probability-weighted averages (Reich et al., 2010) of the tracts, after normalising the parameter maps to MNI space (which was done by applying the warp obtained for the T1-weighted image to the parameter maps). The two approaches are comparable in that voxels with higher tract probability will contribute more strongly to the computed average.

#### 2.2.10. Comparison between individual tracts and probability map-based approach

Firstly, we investigated the anatomical correspondence between the two approaches. We calculated weighted Dice coefficients between individual tract masks (in MNI space) and tract probability maps as done in Hua et al. (2008).

Secondly, we checked whether individually dissected tracts are biased, by omitting damaged parts of the tract. This was done by comparing tract-averaged microstructure of individually dissected tracts to averaged microstructure from the group probability maps. If there was a bias towards the healthy part, the individual measures should indicate less damage than the probability-based measures.

## 3. Results

### 3.1. Comparison of the fibre orientation reconstruction algorithms

#### 3.1.1. Simulation results

*Single fibre population.* All three algorithms could successfully reconstruct the peak of the simulated fibre orientation in almost 100% of substrates with an intracellular volume fraction of at least 20%, which corresponded to an FA value of around .5. The dispersion of successfully reconstructed true peaks consistently lay below .15 (Figure 1). In substrates with intracellular volume fraction below 10%, all algorithms failed to resolve the peak orientation reliably. Here, CSD-based peak detection resulted in a significant number of false positive peaks (up to 25%). On the other hand, the tensor-based approach produced a maximum of 2% false positives. dRL produced less than 0.01% false positives, with the exception of substrates with a single fibre population and the highest simulated fibre content (80% intracellular volume fraction). Inspection of the discrete FOD suggests that this is a truncation artifact of the harmonics that occurred due to the lack of orientation dispersion in the closely packed simulated cylinders. This is something that would only occur in the simulation scenario, as the substrates were simulated with perfectly parallel cylinders, which is a simplification of the axonal arrangement in vivo, where orientation dispersion occurs even in the most systematically packed regions (Mollink et al., 2017). *Crossing fibres.* In simulated substrates with two fibre populations crossing at 90 degrees, by definition, the tensor-based approach only ever reconstructed one of the two peaks. For both dRL and CSD, a minimum total intracellular volume fraction of 40% was needed to consistently detect both true peaks (Figure 1), corresponding to a minimum of 20% per fibre population, which is consistent with what we found in the single fibre population scenario. In substrates with low fibre content, CSD again produced false positives in up to 25% of the voxels, while the false positive rate in the tensor or dRL implementation was negligible. In substrates with intracellular volume fraction above 10%, the dispersion for the truly detected peaks again lay below .15, with the exception of the tensor-based approach in the crossing fibre condition, where the tensor could not reliably represent either of the true fibre orientations.

**Figure 1:**
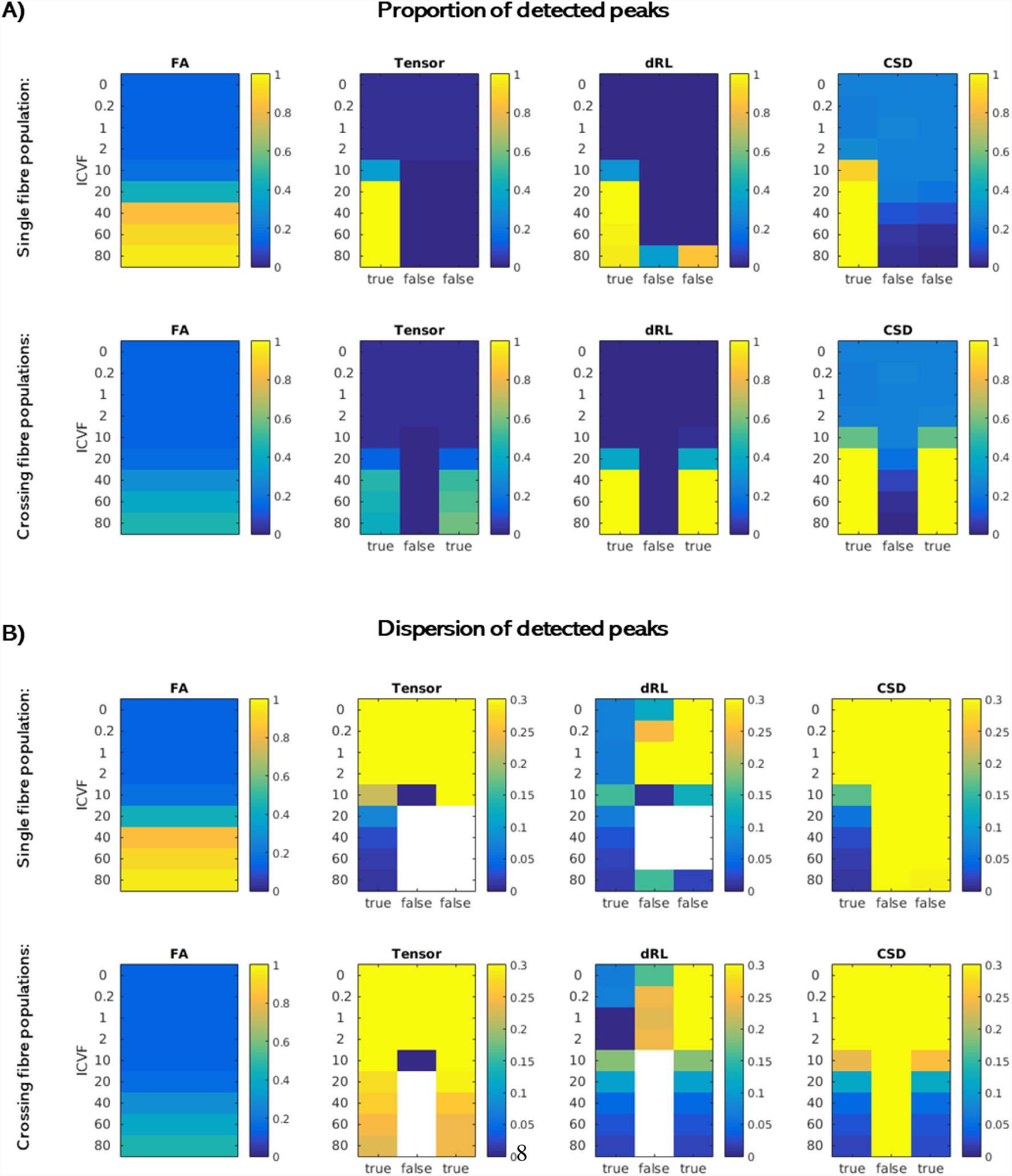
Comparison of the three fibre orientation reconstruction algorithms in simulated data. Simulated substrates varied in their intracellular volume fraction (ICFV). The parallel cylinders in each substrate were aligned with the z-axis (single fibre population), or with the z-as well as the y-axis (crossing fibre populations). **A:** For each approach (tensor-based, dRL and CSD), we calculated the percentage of all voxels within each substrate type for which the ’true’ underlying fibre configuration peak(s) could be detected. As a control, we also calculated this percentage for ’false’ peaks (orthogonal to the true peak(s)). In each case, the left-most plot shows the FA for each substrate type. **B:** Dispersion across all detected peaks of a substrate type was calculated. Dispersion was high across wrongly detected peaks, while for substrates with higher intracellular volume fraction, the true detected peaks by dRL and CSD consistently showed dispersion of < .15.

#### 3.1.2. In vivo results

For each algorithm investigated, we explored its performance within various tissue types: healthy control tissue, normal appearing white matter, white matter lesions that only appear T2-hyperintense, and white matter lesions that also appear T1-hypointense.

The FA and FOD amplitude of continued streamlines in T1-hypointense lesional tissue were slightly lower when comparing to streamlines in lesional tissue without T1-weighted hypointensity. However, overall there were no strong and systematic differences between tissue types (Table 2). On the other hand, with an increasing level of pathology, an increasing number of premature streamline terminations was observed due to the amplitude threshold not being met. This was only the case for the tensor-based approach and dRL (Table 2). In contrast, using CSD, with an increase in pathology, more streamlines were terminated due to the angle criterion. This indicates that while CSD is equally likely to find peaks passing the amplitude threshold in pathological tissue, these peaks are most likely to be spurious.

Counterintuitively, all algorithms produced more streamlines in lesions than in normal appearing white matter, with no significant differences between the two lesional tissue types. This result could be due to an increase of spurious streamlines within lesions, but also due to the preferential localisation of lesions in fibre-rich areas. To check whether the latter might be an explanation for our results, we registered the voxel-wise ’number-of-streamline’ maps from healthy controls to MNI space, and performed voxel-wise correlations between the average number of streamlines and the lesion probability observed in the patients. (This was done for voxels with at least 5% lesion probability, which is the criterion we used to restrict the normal appearing white matter ROI). There were small, but significant correlations, indicating that areas with higher lesion probability in patients also have higher streamline probability in controls (dRL: *r* = .10, *p* < .0001, CSD: *r* = .26, *p* < .0001, tensor: *r* = .18, *p* < .0001).

#### 3.1.3. Considerations of other parameters

As the performance of CSD has been reported to be particularly good at higher b-values (Tournier et al., 2007), we repeated the simulation analyses with b = 2000 s/mm^2^ by increasing the simulated gradient strength, while keeping all other parameters the same. This led to an even higher false positive rate for CSD (Supplementary Figure 2).

The FOD amplitude threshold that was employed for CSD was based on previous work (Jeurissen et al., 2013). Increasing the amplitude threshold from 0.1 to 0.3 eliminated the spurious peaks in the simulated data at b = 1200 s/mm^2^, but did not affect the comparisons in vivo (Supplementary Table 3).

### 3.2. Individual tract segmentation in MS

#### 3.2.1. Evaluation of the tract segmentation methods

Average Dice coefficients of around 80% in healthy controls demonstrate high spatial overlap between manually segmented tracts from two independent operators, indicating robust tract reconstruction protocols (Supplementary Figure 3). To validate the segmentations obtained from the automatic segmentation tool, we also quantified the spatial overlap between automatically segmented tracts and the available manually segmented tracts. The overlap was slightly lower than for the interrater analysis, but still showed an average Dice coefficient of > 60%, in both patients and healthy controls (Supplementary Figure 4). Correlations of tract-specific microstructural metrics extracted from manually vs automatically segmented tracts were high, ranging from *r* = .85 (*p* < .001) to *r* = .98 (*p* < .001) (Supplementary Table 4 and Supplementary Figure 5).

#### 3.2.2. Corticospinal tracts

Overlaying the tract probability map in patients with the lesion probability map (Figure 2 top) confirms that despite the high lesion probability in that region, the individually segmented tracts run through these areas. In all controls and more than 90% of the patients (123/131 for the left and 130/131 for the right), both CSTs could be reconstructed.

**Figure 2:**
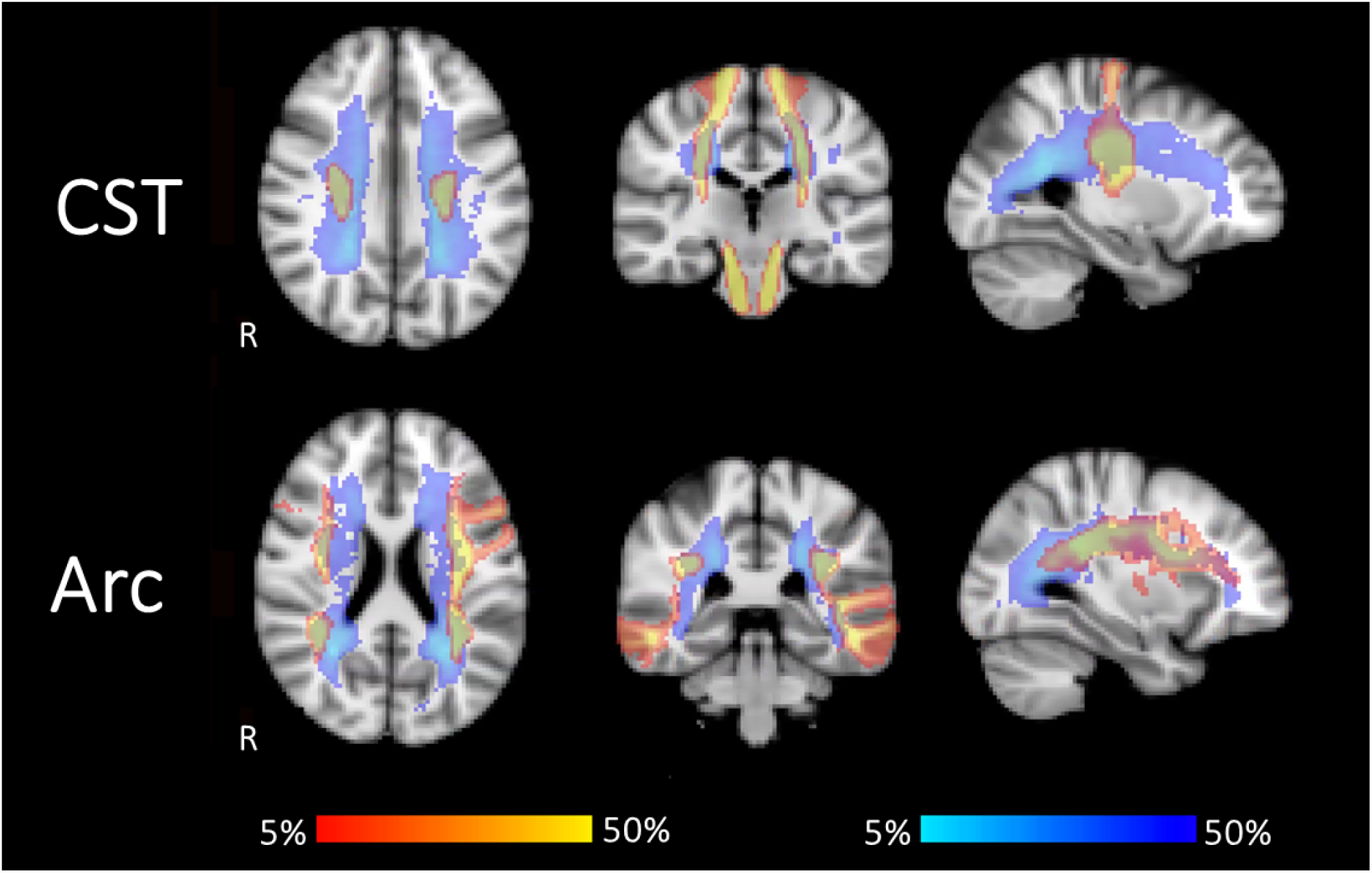
Tract probability. Tract probability maps for patients (red-yellow) are overlaid with lesion probability maps (blue-lightblue). Both are thresholded between 5 and 50 %. Maps for the CST (top) and ARC (bottom) show that areas of high lesion probability overlap with the tracts. This suggests that the investigated tracts go through areas with increased likelihood of lesions, potentially affecting the tracking. R indicates the right hemisphere. **Acronyms:** CST: cortico-spinal tract, ARC: arcuate fasciculus.

#### 3.2.3. Arcuate fasciculi

As evident from overlapping tract and lesion probability maps (Figure 2 bottom), the medial part of the arcuate fasciculus runs is a common location for lesions. In all controls and patients both arcuate fasciculi could be reconstructed, with the exception of the left arcuate fasciculus in 1 patient and the right arcuate fasciculus in 9 patients.

### 3.3. Testing for bias in the individually reconstructed tracts

We extracted average microstructural parameters from individually segmented tracts and from the tract-probability maps that were computed in healthy control data. Average FA and MTR were systematically higher, and average RD was systematically lower in individually reconstructed tracts than with the probability map-based tracts (Table 3). However, the extent to which measures from individual and probability-based tracts differed, was similar in patients and controls, as shown by the non-significant interaction terms (with the exception of MTR in the right CST; Table 3). This indicates that while individually reconstructed tracts systematically differ from the tract probability map, this is the case for both patients and controls, and therefore cannot be attributed to the presence of MS lesions.

**Table 3:** Systematic differences between individually segmented and probability-based tracts. For each of the age-and gender-matched patients (N = 29) and controls (N = 19), mean ± std microstructural metrics (FA, RD and MTR) were extracted from the probability-based (Prob.) as well as for the individually dissected tract mask. To check for systematic differences between individual and probability-based approach, a paired t-test was calculated for each tract, *t* and *p* statistics are reported. To test whether a systematic difference is likely the result of a bias towards the healthy part of the tract in the individual dissections, the difference measure was compared between patients and controls (*t* and *p* values are reported for this interaction.) **Acronyms:** l: left, r: right, CST: cortico-spinal tract, Arc: arcuate fasciculus.

|  | MS |  |  | HC |  |  |  |
| --- | --- | --- | --- | --- | --- | --- | --- |
| FA |  |  |  |  |  |  |  |
| Tract | Prob | Individual | Prob vs Individual | Prob | Individual | Prob vs Individual | Interaction |
| CST l | 0.42 ± 0.03 | 0.50 ± 0.04 | $t = -30.23, p < .01$ | 0.42 ± 0.02 | 0.51 ± 0.02 | $t = -24.35, p < .01$ | $t = -0.03, p = .98$ |
| CST r | 0.42 ± 0.02 | 0.51 ± 0.03 | $t = -54.61, p < .01$ | 0.43 ± 0.02 | 0.52 ± 0.02 | $t = -28.44, p < .01$ | $t = 0.55, p = .58$ |
| Arc l | 0.31 ± 0.02 | 0.41 ± 0.03 | $t = -74.10, p < .01$ | 0.33 ± 0.02 | 0.43 ± 0.02 | $t = -47.95, p < .01$ | $t = -0.66, p = .51$ |
| Arc r | 0.32 ± 0.02 | 0.40 ± 0.03 | $t = -45.85, p < .01$ | 0.34 ± 0.02 | 0.42 ± 0.03 | $t = -21.43, p < .01$ | $t = -0.66, p = .51$ |
| RD (in 10<sup>-3</sup> m<sup>2</sup>/s) |  |  |  |  |  |  |  |
| Tract | Prob | Individual | Prob vs Individual | Prob | Individual | Prob vs Individual | Interaction |
| CST l | 0.66 ± 0.05 | 0.59 ± 0.07 | $t = 13.35, p < .01$ | 0.63 ± 0.04 | 0.55 ± 0.03 | $t = 10.29, p < .01$ | $t = -0.33, p = .75$ |
| CST r | 0.66 ± 0.05 | 0.58 ± 0.04 | $t = 18.78, p < .01$ | 0.64 ± 0.04 | 0.55 ± 0.03 | $t = 12.49, p < .01$ | $t = -1.47, p = .14$ |
| Arc l | 0.71 ± 0.05 | 0.62 ± 0.06 | $t = 44.55, p < .01$ | 0.66 ± 0.02 | 0.57 ± 0.03 | $t = 31.82, p < .01$ | $t = 0.75, p = .46$ |
| Arc r | 0.69 ± 0.05 | 0.61 ± 0.06 | $t = 30.85, p < .01$ | 0.64 ± 0.02 | 0.57 ± 0.03 | $t = 21.80, p < .01$ | $t = -0.05, p = .96$ |
| MTR |  |  |  |  |  |  |  |
| Tract | Prob | Individual | Prob vs Individual | Prob | Individual | Prob vs Individual | Interaction |
| CST l | 0.40 ± 0.01 | 0.42 ± 0.02 | $t = -23.46, p < .01$ | 0.40 ± 0.01 | 0.42 ± 0.01 | $-16.07, p < .01$ | $t = 0.92, p = .36$ |
| CST r | 0.39 ± 0.01 | 0.42 ± 0.01 | $t = -38.89, p < .01$ | 0.40 ± 0.01 | 0.43 ± 0.01 | $-22.42, p < .01$ | $t = 2.49, p = .01$ |
| Arc l | 0.39 ± 0.02 | 0.41 ± 0.02 | $t = -50.04, p < .01$ | 0.40 ± 0.01 | 0.42 ± 0.01 | $-46.33, p < .01$ | $t = 1.18, p = .24$ |
| Arc r | 0.39 ± 0.02 | 0.41 ± 0.02 | $t = -34.77, p < .01$ | 0.40 ± 0.02 | 0.42 ± 0.02 | $-24.59, p < .01$ | $t = 1.92, p = .06$ |

### 3.4. Anatomical correspondence between individually segmented and probability-based tracts

We quantified the anatomical overlap between the individually dissected tracts and the group probability map with a probability weighted overlap score (see Hua et al. (2008)). In both patients and controls, spatial overlap varied between tracts, with median scores of around 55% for the CST and of around 35% for the arcuate fasciculus (Figure 3). The individual variability suggested that for some patients only 10% of voxels were shared between individual tract mask and probability mask, while for others it was 75%.

**Figure 3:**
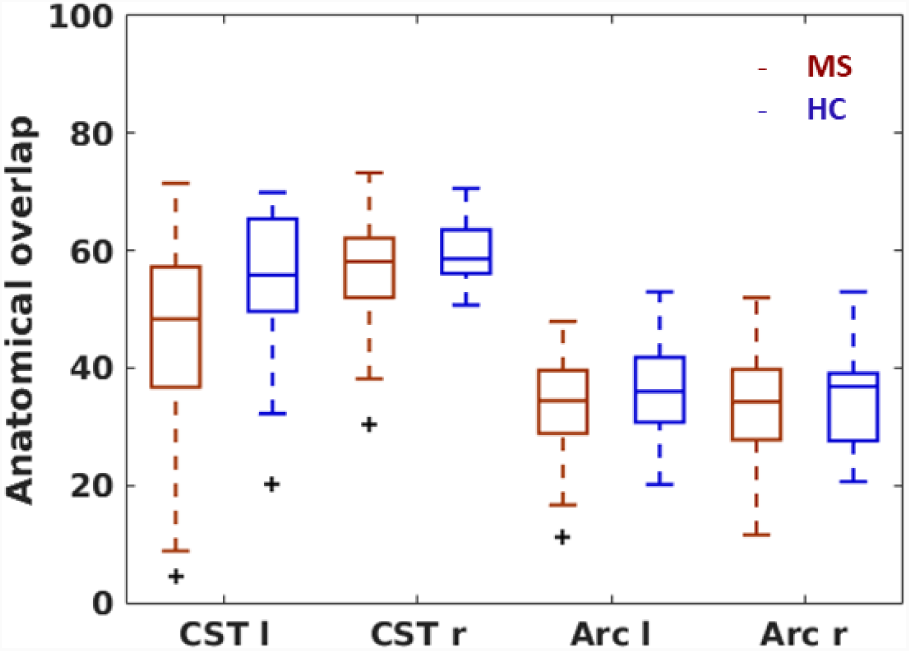
Anatomical overlap. An overlap score between the tract probability map and the individual tract mask (registered to MNI space) were calculated. Each time, boxplots for patients (bright) and controls (dark) are presented for each tract. Mean ± std of the weighted Dice coefficients (converted to %) are 45±14 (MS) and 55±13 (HC) for the left CST, 57±8 (MS) and 60±6 (HC) for the right CST, 45±8 (MS) and 36±8 (HC) for the left arcuate fasciculus, and 32±9 (MS) and 35±8 (HC) for the right arcuate fasciculus. **Acronyms:** l: left, r: right, CST: cortico-spinal tract, ARC: arcuate fasciculus.

### 3.5. Correspondence in tract-specific microstructure of individual and probability-based approach

We also assessed the agreement between tract-based estimates of microstructural damage between the individual and the probability-map approach. Correlations were very high for the arcuate fasciculus, ranging from *r* = .87 (*p* < .001) to *r* = .97 (*p* < .001), and lower for CST, ranging from *r* = .49 (*p* < .001) to *r* = .91 (*p* < .001). Results were similar for patients and controls (Figure 4).

**Figure 4:**
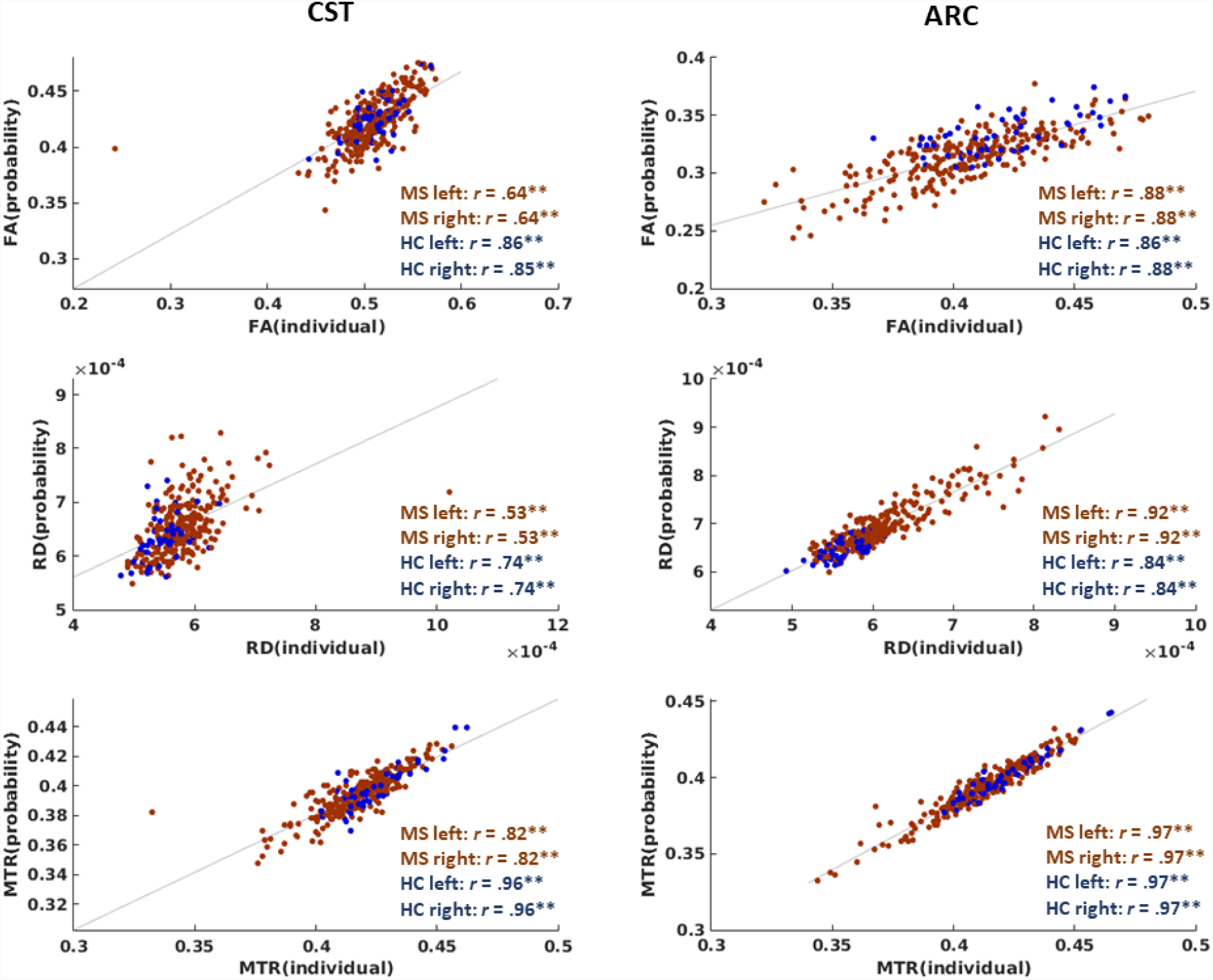
Correlations between tract-specific microstructure from individually segmented and probability-based tracts. For each microstructural metric (FA, RD and MTR; rows) and tract (columns), the correlation between individual approach and probability-based approach is shown. Data were collapsed across hemispheres for plotting. Pearson correlation coefficients are provided for each group and hemisphere separately. MS data points are plotted in red, HC in blue. The line of best fit across all data points is shown. \*\**p* < .0001. **Acronyms:** CST: cortico-spinal tract, ARC: arcuate fasciculus.

## 4. Discussion

In this work we revisited the challenges of tracking through pathological tissue. We showed that the presence of MS lesions affects fibre orientation reconstruction algorithms differentially: during tensor-and dRL-based tracking, streamlines are more likely to stop, while CSD is more likely to produce spurious streamlines. Even though the CST and arcuate fasciculi run through regions that are frequently affected by MS lesions, we were able to successfully perform individual tract segmentation in a large patient cohort. The segmented tracts did not show a systematic bias in the estimation of microstructural health, when compared to a frequently employed approach based on tract probability maps. While tract-averaged microstructural measures showed medium to high correlations between the two approaches, the anatomical correspondence was limited, highlighting the potential benefits of individual tract reconstruction.

### 4.1. MS pathology affects fibre orientation reconstruction algorithms differentially

To compare fibre orientation reconstruction algorithms under controlled conditions, we simulated tissue substrates that varied in the number of fibres they contained, providing a simplified model of damage in MS. Lesions are characterised by a variety of pathological processes, with fibre loss likely being the change that is the most significant for tractography. Fibre loss could lower the amplitude of the FOD peaks, which are followed during streamlining. At the same time, the associated increase in extracellular water is likely to impact on the diffusion profile, which could further complicate the identification of peaks. Indeed, in our simulated substrates with intra-cellular volume fractions below 10%, none of the algorithms could reliably identify the true underlying fibre orientations. With intracellular volume fractions above 10%, all three algorithms were consistently successful, with the exception of the tensor-based approach under crossing fibre conditions due to its inherent constraint that it can only ever reconstruct one peak. The main difference between CSD and dRL, in both the single and crossing fibre scenarios, was the false positive rate. With the specified FOD amplitude thresholds, only CSD, but not dRL, produced a substantial amount of spurious peaks in substrates with lower fibre content and low FA. This could be a result of a mismatch between responses in high-FA calibration tissue and the target responses (Parker et al., 2013b).

The findings from the simulations were largely supported by in vivo data from MS lesions. DRL seemed to be more conservative than CSD, with an increase in streamline termination in lesions due to low FOD amplitudes. Premature termination was most pronounced in T1-hypointense lesions, presumably due to the particularly high fibre loss found in this lesion type (Sahraian et al., 2010). dRL has previously been shown to perform worse in tissue with crossing fibres and low FA (Parker et al., 2013b), which could be the reason for the increased streamline terminations in lesion. Even though CSD may be able to resolve fibres better under these conditions (Parker et al., 2013b), our results suggest that it may be more sensitive to the increased isotropic diffusion in lesions than dRL, producing false positive peaks and consequently spurious streamlines. Even though CSD’s performance at resolving crossing fibres has been reported to be better for data obtained at high b-values (Jeurissen et al., 2013), in our case, increasing the b-value during the simulation led to an even higher false positive rate in CSD. It is possible that the FOD threshold for peak detection, which we based on previous studies (Jeurissen et al., 2013; Parker et al., 2013b) is not universally optimal for all types of data, SNRs, diffusion-weightings etc. In our case, increasing the FOD threshold from 0.1 to 0.3 reduced the false positives rate during the simulations, but did not change the behaviour of the algorithm in vivo. It is likely that optimal tractography parameters for simulated data are not the same optimal parameters for in vivo data, as simulations are often simplified, e.g. in our case the fibres in the simulated substrates lacked orientation dispersion. Systematic and thorough parameter tuning in pathological in vivo data could potentially help to optimise the use of CSD in MS lesions, but was out of scope of this study.

Choosing the algorithm for tract segmentation in vivo, we considered two things. Firstly, dRL seems to be the more conservative algorithm, which leads to streamline termination in lesions with high fibre loss, potentially producing tract reconstructions that at least partially omit lesions. CSD produces more streamlines in lesions, but a considerable number of them may be spurious, which also hinders the correct reconstruction of tracts that pass through lesions. The second consideration was theoretical. In contrast to dRL, CSD requires the response function used for the deconvolution to be calibrated from voxels with single underlying fibre populations, which are generally identified through their high FA. In MS, high FA voxels are likely voxels with preserved white matter integrity. However, the single fibre response function is then applied to the lesional tissue, which could lead to problems with FOD reconstruction and tracking in lesional tissue. Estimating a separate ’pathological’ single fibre response function from the average response in lesioned voxels is unlikely a sensible alternative, as lesional tissue is highly heterogeneous. In comparison to CSD, dRL is more robust to mis-calibrations of the single fibre response function (Parker et al., 2013b). For this reason, for the purpose of this paper, we performed the in vivo whole-brain tractography in MS using dRL.

### 4.2. Fibre tracking in MS allows to reconstruct tracts that go through areas with high lesion probability

Even though the CSTs and arcuate fasciculi pass through white matter regions that are frequently affected by lesions, we successfully segmented these tracts in individual brains from a large cohort of MS patients. A previously introduced automated tract extraction algorithm made this a feasible endeavour (Parker et al., 2013a). Unlike the automated approach using tract probability maps (Reich et al., 2010), this method learns the shape and approximate location of a tract from training data and uses the resulting model to identify and segment tracts from wholebrain tractograms, obtained with the same tractography pipeline as in the training data. We found that the automated approach produces tracts that are realistic with regard to their shapes and anatomical location and that show high spatial overlap with manually segmented tracts. In all our spatial agreement analyses, for some individuals the spatial overlap between tracts obtained through different methods was low. This suggests that even when standard manual segmentation protocols are robust, the process is not completely objective and there is still room for error when drawing anatomically-defined waypoints. Our automatic segmentation method provides a feasible tool for individual tract segmentation in a large number of patients.

An obvious reason to be careful about individual tract segmentation in MS, even when done in an automated manner, is that if lesions or otherwise damaged parts of the tract are left out by the tracking algorithm, then there would be a systematic bias in tract-averaged microstructural parameters towards values from healthier tissue. We found that individually segmented tracts show higher FA, lower RD and higher MTR than when employing probability-based tract masks, suggesting the opposite, so more intact microstructure in individually segmented tracts. This is not a surprising result, as the tractography algorithm will preferentially produce streamlines in voxels with high white matter and fibre density. In contrast, the probability-map approach considers all voxels in the probability map, independent on the underlying white matter in an individual patient. However, importantly, the reported systematic difference between individual and probability-map based tracts we found was comparable between patients and controls. This finding suggests that there is no additional disease-related bias in the individual tract segmentations. While the simulation and tissue comparison results suggest that the fibre orientation reconstruction algorithms could further be optimised for tracking through lesions, our tract segmentation results suggest that dRL may be a promising method for tractography in MS.

#### 4.2.1. The anatomical correspondence of individual and probability-based approach is modest

Previous studies have suggested using a probability-based approach of investigating microstructure in specific white matter tracts in MS. Here, we showed that the spatial overlap between tract probability maps obtained from healthy controls, and individually segmented tracts of individuals is modest, with averages ranging from 35% for the right arcuate fasciculus to 60% for the right CST, with similar results for patients and controls. The stronger agreement for the CST suggests that there may be more individual variability in tract shape and localisation in the arcuate fasciculus. In both cases, the variability in individual tracts is likely to be partially also a result of the normalisation of individual brains to MNI space, which does not necessarily align tracts within the homogeneously appearing white matter.

Other methodological factors are likely to play a role as well. Even though the anatomical correspondence between individual and probability-based approach was lower than the correspondence between manual and automatic segmentations and the correspondence between manual segmentations of different operators, latter methodological differences also affected the tract anatomy. Without ground truth data available, it is difficult to conclude which methods give the most accurate tract segmentations (Schilling et al., 2019).

#### 4.2.2. The correlation of tract-specific microstructure in individually segmented vs probability-based tracts is high

Despite the difference in spatial location and systematic difference in average microstructural parameters, the correlation of the microstructural information between the individual and probability-map based approaches was high, particularly for the arcuate fasciculus. Correlations between microstructural parameters from individually segmented tracts vs atlas-based estimations have previously been reported to be tract-dependent (Reich et al., 2010). Some of our correlations were also lower, e.g. only 25% between the two approaches were shared in RD measures of the right CST. Low correlations for tensor-based measures, such as RD, could be influenced by their sensitivity to the macroscopic tract shape and individual differences in tract morphometry (De Santis et al., 2014). This hypothesis is corroborated by the result that the correlation between individual and probability-based approach in the CST was highest for MTR, a microstructural metric that does not show this sensitivity to tract morphometry.

Individual tract segmentation may not bring large benefits compared to the probability-based approach, when average microstructural parameters of a tract as a whole are of interest. This can be the case for studies looking at global effects within specific tracts (e.g. Bonzano et al. (2014)). Here, averaging microstructural measures should be less noisy and provide more statistical power and spatial accuracy than voxel-based approaches. However, the advantage of individual tract segmentation is that it does not limit the analysis to tract-specific average measures. Other features of the tract can also be studied, such as tract shape, tract-specific atrophy (Wang et al., 2018) or the spatial variation along a tract (Jones et al., 2005; Yeatman et al., 2012). Very precise anatomical -rather than probabilistic assignment -of lesions to specific white matter specific tracts could also open doors for clinical questions, for example on how the appearance or disappearance of lesions in specific tracts is associated with the presence or resolution of specific symptoms or with potential secondary damage, such as pathology in the cortical regions that are directly connected by a tract.

#### 4.2.3. Limitations and future directions

Under the conditions of this study, CSD produced more spurious peaks than the other algorithms. However, this is not to say that the algorithm does not have benefits which could aid tractography in MS patients. For example, CSD has previously been shown to outperform dRL when resolving crossing fibres under low FA conditions (Parker et al., 2013b). We only assessed the performance of the algorithms in the context of tractography, where not all FOD peaks are reconstructed and considered simultaneously. Instead, at each point along the streamlining process, the closest detected peak to the current streamline direction is followed, assuming that the tangent to the streamline minimally subtends the best estimate of fibre orientation. To mimic the tractography process in our simulation, the 45 degree angle threshold employed during streamlining was also employed when assessing the proportion of successfully detected peaks. This is a comparatively lenient threshold for assessing peak detection accuracy, with an additional measure of dispersion confirming orientational agreement between the detected peaks. While under these specific conditions, CSD produced a large number of false positives, at this point we are not in a position to comment of the suitability of CSD for other types of analyses, such as fixel-based analysis (Raffelt et al., 2015).

We employed standard tractography protocols that have been optimized in healthy tissue, and that are compatible with clinically feasible diffusion MRI acquisitions. It is likely that optimising tractography for MS could benefit from fine-tuning of some of the parameters for pathological tissue, and also from data with higher angular resolution and multiple diffusion weightings. For example, the sensitivity of CSD to the increase in isotropic diffusion in lesions may be partly counteracted using multi-shell -CSD, which considers several tissue compartments (Jeurissen et al., 2014). Transitioning to multi-shell acquisitions could allow the benefits of such advanced deconvolution methods to be explored.

Simulations produced a simplified version of pathology, and other factors such as permeability, could also be considered to make the scenario more realistic (e.g. Nedjati-Gilani et al. (2017)). Ideally, also fibre orientation dispersion is introduced, which is currently not possible with the Camino software package used in the current study.

The probability-based approach depends on inter-subject co-registration of the images by normalising them to a common reference space. To do this, we chose T1-weighted based co-registration, as this is most successful for aligning pathological brains (Avants et al., 2008), and to make our results comparable to previous studies (Reich et al., 2010). However, in future studies, this normalisation step could be further improved, e.g. in the form of multi-modal registration by white matter parameteric maps, such as FA maps.

#### 4.2.4. Conclusions

Accurate anatomical assignment of damage to specific white matter tracts is of clinical interest. In MS research, tractography-based anatomical segmentation in individual patients is rarely performed, with probability-based approaches often being the method of choice to avoid potential effects of MS lesions on tractography algorithms. We show that MS pathology indeed affects the fibre orientation reconstruction that is done during tractography, with different algorithms being affected differently -lesions led to an increase in streamline interruptions with dRL, and an increased number of spurious streamlines with CSD. Nevertheless, fiber tracking through MS lesions is possible, and can be used to even reconstruct tracts that go through areas with high lesion probability. The resulting tracts do not seem to systematically omit lesional tissue. A problem with tractography in general is that the anatomical localisation of the tract does depend on the specific method used to segment individual tracts. The anatomical overlap between individual tracts and tract probability methods is quite low, however, this may not be an issue if the aim of the research is to obtain tract-averaged microstructural parameters. For studying tract shapes, tract-specific atrophy and along-tract profiles, individually segmenting tracts is necessary. We showed that this is feasible, even in large scale clinical studies. Further improvements of the algorithms to maximize anatomical accuracy could lead to powerful tools to investigate prognostic and diagnostic markers.

## 4.3. Acknowledgements

The study was funded by a research grant of the MS Society UK. DKJ is supported by a Wellcome Trust Investigator Award (096646/Z/11/Z) and a Wellcome Trust Strategic Award (104943/Z/14/Z). EP is funded by the Wellcome Trust.

The authors would like to thank Matt Hall for his input on the data simulations.

## Supplementary material

### 1.1.1 Supplementary methods: Data simulation parameters

We simulated diffusion data using Camino’s datasynth (Hall and Alexander, 2009). We simulated data using a virtual pulse sequence, comparable to the in vivo sequence (twice-refocussed spin echo (TRSE), TE = 94.5 ms, b-value = 1200s/mm^2^, δ1: 11.2 ms, onset δ1: 15.2 ms, δ2: 17.8 ms, onset δ2: 31.7 ms, δ3: 17.8 ms, onset δ4: 75.3 ms, gradient amplitude: 40 mT), with 40 uniformly distributed gradient directions (Camino 40), with 6 non-diffusion weighted images at the beginning.

We simulated substrates with parallel cylinders, with radius drawn from a gamma distribution with a shape parameter of 2 and scale parameter of 5 × 10^−^7, which corresponds to a mean radius of 1 *µ*m and standard deviation of 0.7 *µ*m. The parameters used in the simulation were: 500000 walkers (numbers of spins simulated), uniformely distributed across the substrate, cylinder permeability 0, tmax = 5000. We simulated various substrates (cubes of the length 50 × 10^−^5 m), that only differed in their cylinder density (0 to 40000 cylinders placed in substrate), yielding intracellular volume fractions of about 0 −80%. For each substrate type, we simulated 20 different substrates (by choosing different seed values), and for each substrate we simulated 100 voxels that only differed in their noise. The SNR chosen for simulation was 20:1, which was based of a conservative estimate derived from our in vivo data (see below). In order to simulate crossing fibres, we ran the exact same simulation, but rotating the acquisition scheme around the x-axis by 90 degrees.

### 1.1.2. Supplementary methods: Estimation of SNR from in vivo data

To estimate the SNR of our in vivo diffusion sequence, we used the SNR_mult_ approach described by Dietrich et al. (2007). This relies on calculating the ratio between the mean of the signal divided by the standard deviation of the underlying (Gaussian) noise. To do this, the six non diffusion-weighted images were taken as repeated acquisitions. An SNR map was calculated for 29 MS patients, followed by registration of each map to MNI space. We averaged the SNR maps across participants. Thresholding the SNR maps at 20 showed that this value was exceeded across the whole brain, with the exception of the putamen. From this, we concluded that a SNR of 20 was a conservative estimate. (Note that in the calculation, the noise was assumed Gaussian, which is the case for SNR > 3 (Gudbjartsson and Patz, 1995). However, the SNR was estimated based on non-diffusion weighted images, and the diffusion-weighted images have lower signal. To check that the signal attenuation was not more than 1/7 (which would cause the SNR in the attenuated images to fall below 20), we visualised in a few data sets the maximum signal attenuation, and in the vast majority of voxels this was not the case.)

### 1.1.3. Supplementary methods: Fibre orientation algorithm implementation

The diffusion tensor was derived by robust non-linear least squares fitting (Chang et al., 2005), using ExploreDTI v.4.8.3.

The constrained spherical convolution (CSD) Tournier et al. (2007) was implemented using in-house scripts. The single fibre response function was calculated from voxels with an FA > .8, as done in previous work (Tournier et al., 2004). For the simulated data, the CSD response function was estimated from voxels with FA > .8 in the single fibre population data set. Spherical harmonics were resolved up to the 6th order. During the tractography process, we employed an FOD amplitude threshold of > 0.1 that has been previously optimised (Jeurissen et al., 2013).

The modied damped Richardson-Lucy algorithm (dRL) was implemented with a fibre response shape parameter of α = 1.5*x*10 −3*mm*^2^/*s* according to Dell’Acqua et al. (2010) using in-house scripts. Note that we fitted harmonics up to the 8th order to the discrete dRL estimates, to increase computational efficiency, while allowing to track along all potential directions rather than only along the discretely estimated directions. The FOD amplitude threshold was set to > 0.05 (Parker et al., 2013b).

### 1.1.4. Supplementary methods: Segmentation protocols

#### CST

To segment the CSTs, AND gates were placed in the primary motor cortex (identified on the T1-weighted image) and in the brain stem (identified as the blue colour of the pons in the anterior part of the brain stem in the axial slice of a first eigenvector-colored FA image (Pajevic and Pierpaoli, 1999)). This protocol is comparable to Mole et al. (2016). Left and right CST were segmented separately.

#### Arcuate

Left and right arcuate fasciculi were segmented separately. To do so, each time first a coronal slice of a first eigenvector-colored FA image was identified, in which the posterior commissure was visible. Then a SEED gate was drawn in the arcuate fasciculus, identified as a green triangle lateral to the corpus callosum. Additionally, an AND gate was drawn where the arcuate fasciculus bends, identified as the blue / purple appearing ipsilateral to the SEED gate on an axial slice at the height of the posterior commissure.

#### 1.1.5. Calculation of spatial agreement / Dice coefficient

We calculated spatial overlap Dice coefficient scores (see also Zijdenbos et al. (1994)) as follows: First the tracts were exported to binary NIfTI files, then the number of voxels for each operator’s tract and the number of voxels overlapping in both operators’ tracts were counted using the AFNI function *3DOverlap* (Cox, 1996). Finally, Dice coefficients were calculated using the equation: 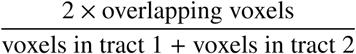, and converted to %.

**Table S2:**
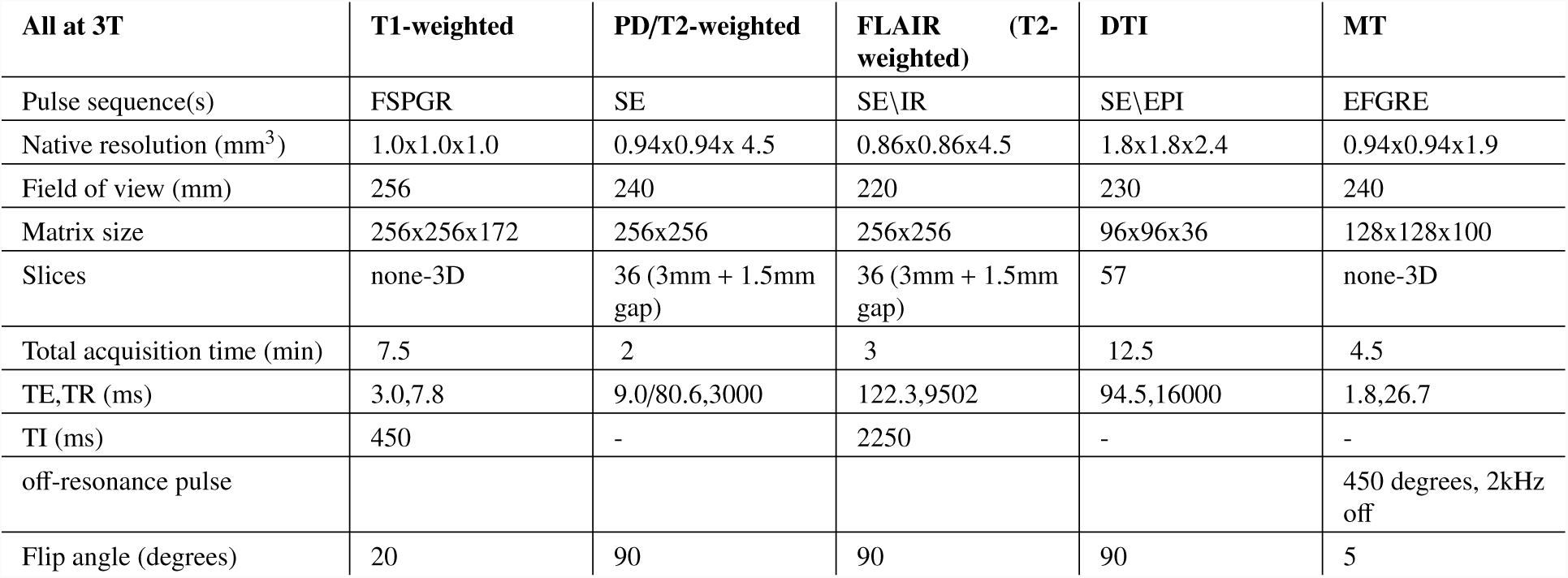
Scan parameters. All sequences were acquired at 3T. For each of the sequences, the main acquisition parameters are provided. **Acronyms:** FLAIR = fluid-attenuated inversion recovery, FSPGR = fast spoiled gradient echo, SE = spin-echo, IR = inversion recovery, EPI = echo-planar imaging, EFGRE = enhanced fast gradient echo, TE = echo time, TR = repetition time, TI = inversion time.

**Table S3:**
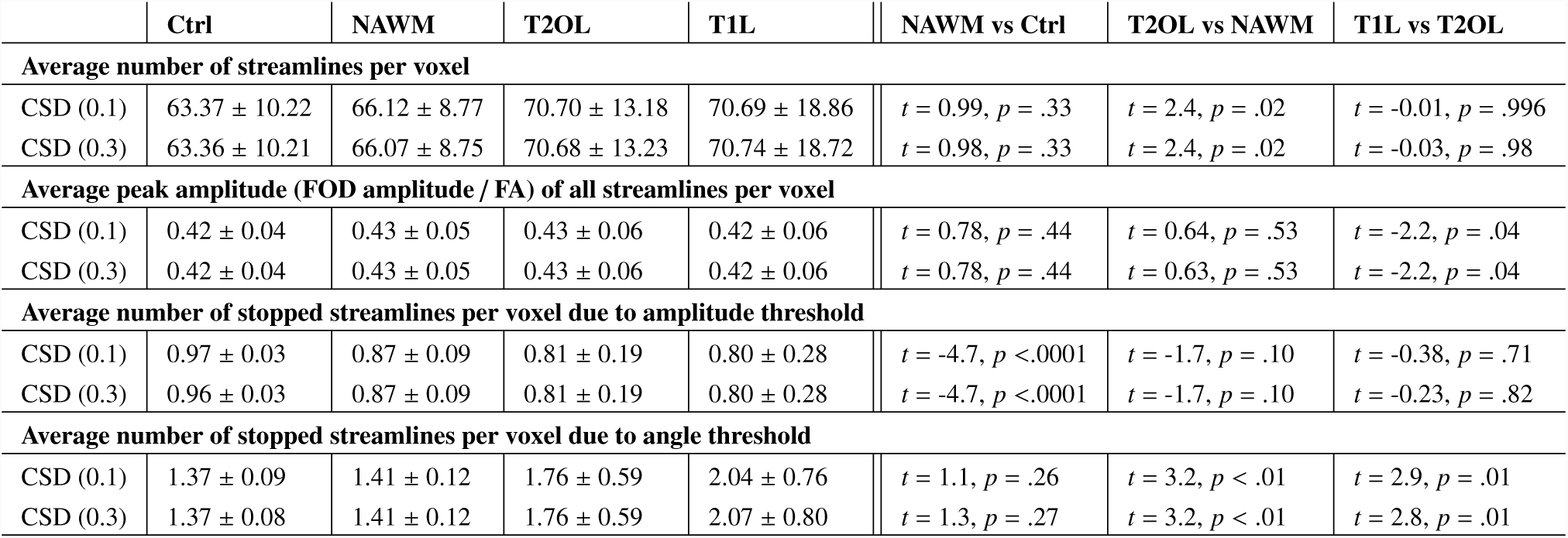
Comparison of FOD threshold for CSD in vivo. As done before, for both CSD FOD thresholds (0.1 and 0.3) and each tissue type (Ctrl, NAWM, T2L, T1l), we calculated voxel-wise averages of the following parameters: the number of streamlines found in a voxel, the average FA / FOD amplitude across all streamlines found in a voxel, the number of streamlining processes stopped due to the amplitude threshold in a voxel, and the number of streamlining processes stopped due to the angle threshold in a voxel. The mean ± std of these measures across healthy controls (Ctrl; N = 19) and patients (NAWM, T2L, T1L; N = 29) are reported. The values across different tissues types were statistically compared (unpaired t-test between Ctrl and NAWM tissue; paired t-tests for T2OL vs NAWM, and T1L vs T2OL).

**Table S4:** Correlations between tract-specific average microstructural metrics for automated vs manual tract dissection. For each metric and tract and each group separately, the sample size (N) and the Pearson correlation coefficient and the corresponding *p*-values are reported. From Figure 5 it is evident that some low correlations may be caused by outliers. For these correlations, bivariate outliers were excluded (based on (Rousselet and Pernet, 2012)) and outlier-robust correlations and corresponding *p*-values are also reported. Note that correlations were not systematically lower in patients, even though the tracts from the healthy controls were used to create the model for automated tractography. **Acronyms:** l: left, r: right, CST: cortico-spinal tract, Arc: arcuate fasciculus, FA = fractional anisotropy, RD = radial diffusivity, MTR = magnetisation transfer ratio.

|  | FA |  |  |  |  |  |
| --- | --- | --- | --- | --- | --- | --- |
|  | N MS | N HC | MS | MS robust | HC | HC robust |
| CST l | 28 | 19 | $r = .63, p < .001$ | $r = .96, p < .001$ | $r = .86, p < .001$ | $r = .86, p < .001$ |
| CST r | 29 | 19 | $r = .51, p = 0.01$ | $r = .85, p < .001$ | $r = .91, p < .001$ | $r = .91, p < .001$ |
| Arc l | 29 | 19 | $r = .61, p < .001$ | $r = .97, p < .001$ | $r = .49, p = 0.03$ | $r = .98, p < .001$ |
| Arc r | 26 | 18 | $r = .56, p < .001$ | $r = .94, p < .001$ | $r = .68, p < .001$ | $r = .88, p < .001$ |
|  | RD |  |  |  |  |  |
|  | N MS | N HC | MS | MS robust | HC | HC robust |
| CST l | 28 | 19 | $r = .51, p = .01$ | $r = .95, p < .001$ | $r = .91, p < .001$ | $r = .93, p < .001$ |
| CST r | 29 | 19 | $r = .52, p < .001$ | $r = .89, p < .001$ | $r = .86, p < .001$ | $r = .91, p < .001$ |
| Arc l | 29 | 19 | $r = .75, p < .001$ | $r = .96, p < .001$ | $r = .57, p = 0.01$ | $r = .88, p < .001$ |
| Arc r | 26 | 18 | $r = .76, p < .001$ | $r = .98, p < .001$ | $r = .72, p < .001$ | $r = .96, p < .001$ |
|  | MTR |  |  |  |  |  |
|  | N MS | N HC | MS | MS robust | HC | HC robust |
| CST l | 28 | 19 | $r = .78, p < .001$ | $r = .98, p < .001$ | $r = .98, p < .001$ | $r = .96, p < .001$ |
| CST r | 29 | 19 | $r = .85, p < .001$ | $r = .85, p < .001$ | $r = .97, p < .001$ | $r = .93, p < .001$ |
| Arc l | 29 | 19 | $r = .66, p < .001$ | $r = .99, p < .001$ | $r = .92, p < .001$ | $r = .85, p < .001$ |
| Arc r | 26 | 18 | $r = .80, p < .001$ | $r = .96, p < .001$ | $r = .88, p < .001$ | $r = .88, p < .001$ |

**Figure S2:**
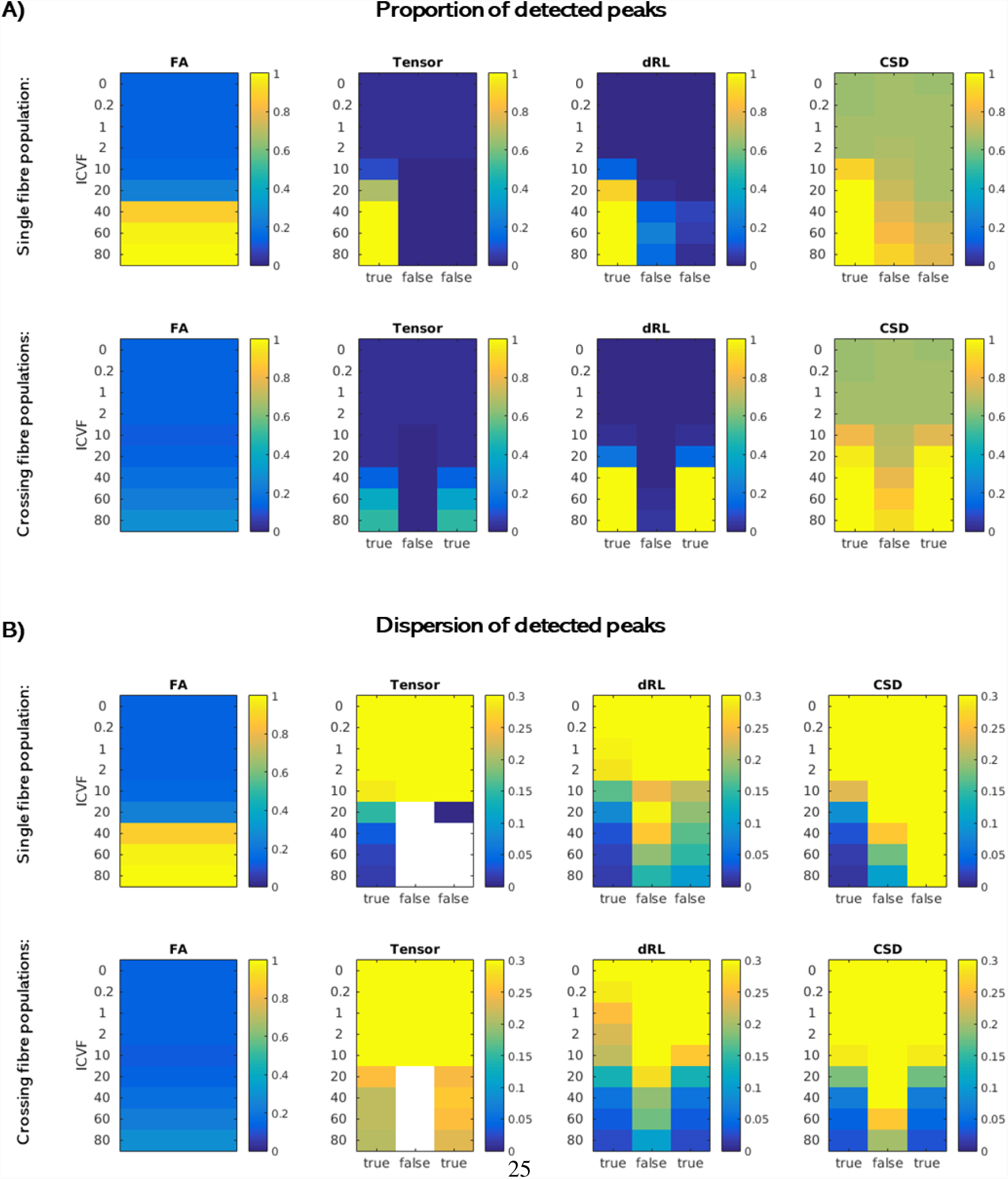
Comparison of the three fibre orientation reconstruction algorithms in simulated data with b *=* 2000 s*/*mm^2^. Simulated substrates varied in their intracellular volume fraction (ICFV). The true fibre orientation of the parallel cylinders in each substrate was along the z-axis (single fibre population), or along the z-as well as the y-axis (crossing fibre populations). **A:** For each approach (tensor-based, dRL and CSD), we calculated the percentage of all voxels within each substrate type for which the ’true’ underlying fibre configuration peak(s) could be detected. As a control, we also calculated this percentage for ’false’ peaks (orthogonal to the true peak(s)). In each case, the left-most plot shows the FA for each substrate type. **B:** Dispersion across all detected peaks of a substrate type was calculated.

**Figure S3:**
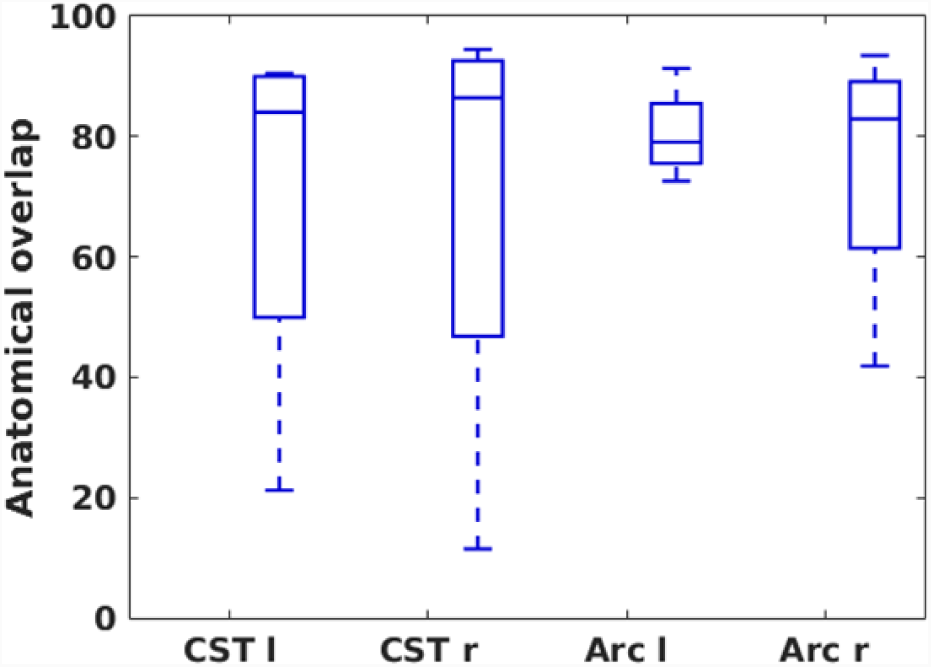
Inter-operator anatomical agreement. Spatial Dice coefficients were computed to quantify the overlap between segmented tracts from two independent operators in %. This was done for tracts from five healthy controls. Each time, boxplots are presented for each tract. **Acronyms:** l: left, r: right, CST: cortico-spinal tract, ARC: arcuate fasciculus.

**Figure S4:**
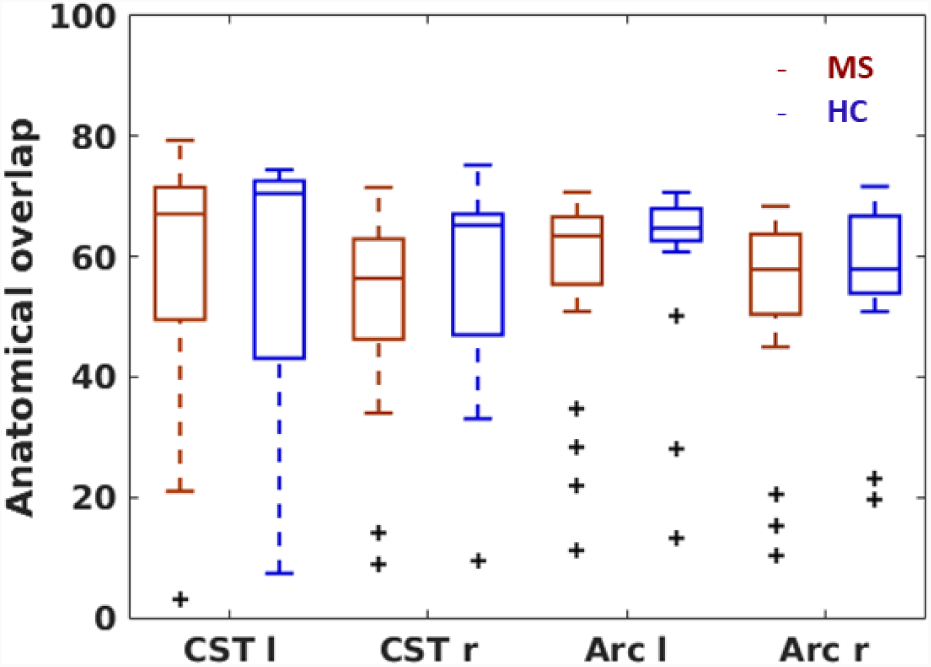
Anatomical overlap between manually and automatically segmented tracts. Spatial Dice coefficients were computed to quantify the overlap between manually and automatically segmented tracts in %. Each time, boxplots for patients (red) and controls (blue) are presented. Outliers are indicated with black crosses **Acronyms:** l: left, r: right, CST: cortico-spinal tract, ARC: arcuate fasciculus.

**Figure S5:**
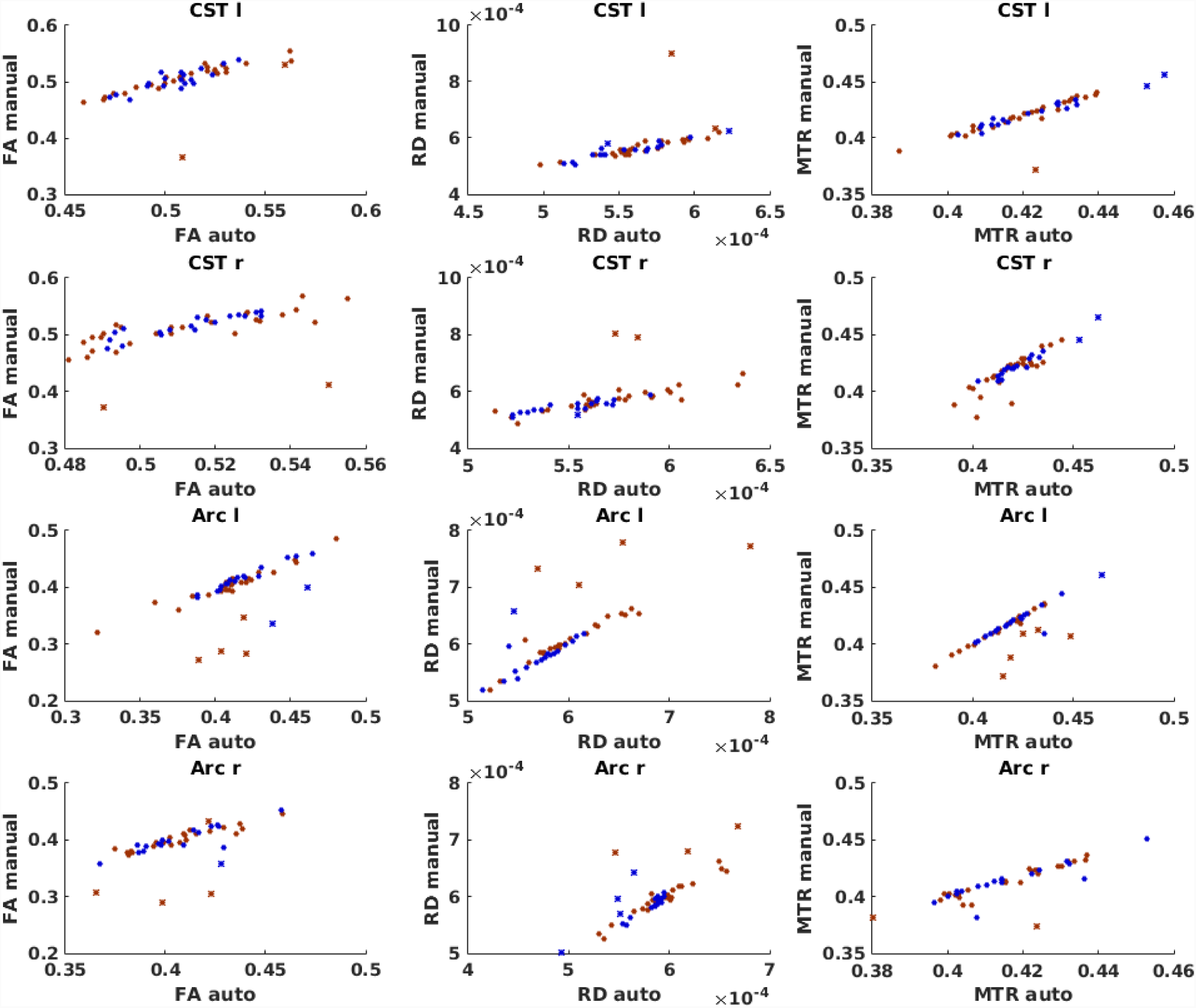
Correlations between tract-specific metrics for automated vs manual dissection. For each metric (rows) and tract (columns), the scatterplot for the correlation between automated and manual approach is shown. MS patients are represented by red dots, and healthy controls by blue dots. In each case, identified bivariate outliers are represented with the asterix symbol. **Acronyms:** l: left, r: right, CST: cortico-spinal tract, ARC: arcuate fasciculus, FA = fractional anisotropy, RD = radial diffusivity (in 10^−3 m2^/s), MTR = magnetisation transfer ratio.

